# T cell memory alters pulmonary inflammatory responses to cecal ligation and puncture

**DOI:** 10.1101/2025.05.15.654338

**Authors:** Mariana R. Brewer, Clifford S. Deutschman, Matthew D. Taylor

**Author notes:** **Corresponding Author:** Mariana R. Brewer Northwell, New Hyde Park, NY Cohen Children’s Medical Center Division of Neonatology, 269-01 76^th^ Avenue, Suite 344 New Hyde Park, NY 11040, Work. **Conflicts of Interest and Source of Funding:** Mariana Brewer received funding from Cohen Children’s Medical Center Intramural Mentored Pediatric Service Line Research Grant for Early Career Investigators. Matthew Taylor receives funding from an NIGMS R35 award (GM157197). Clifford Deutschman is a consultant to Enlivex Inc, Jerusalem Israel, is the co-editor of a textbook published by Elsevier Inc for which he receives royalties and receives a yearly stipend as Scientific Editor of *Critical Care Medicine*. Drs. Deutschman and Taylor have received doses of angiotensin-II (Giapreza) from La Jolla Pharmaceuticals for use in research. These potential conflicts do not directly relate to the findings detailed in this paper.

## Abstract

To date, murine sepsis models have failed to recapitulate human acute respiratory distress syndrome, one of the leading complications of human sepsis. We set out to determine if preexisting T cell memory, which is common in human adults and lacking in laboratory mice, could contribute to lung inflammation in the cecal ligation and puncture (CLP) model of sepsis. After administering an anti-CD3ε activating antibody to C57Bl/6 mice to induce a T cell memory repertoire, we compared the pulmonary immune response to CLP in these “Immune-Educated mice” to responses observed in Uneducated control animals. Compared to Uneducated mice, 24 hours after CLP, Immune-Educated mice had higher alveolar inflammatory cytokine and chemokine concentrations and more pulmonary interstitial macrophages. After 48 hours, the proportion of effector CD4 T cells that produced interferon-gamma was greater in Immune-Educated mice. After 72 hours, there were more alveolar macrophages in the lungs of Educated mice. Separately, we performed adoptive transfer of memory CD4 and CD8 T cells from immunized C57Bl/6J to B6.SJL mice and IFNγ blockade at the time of CLP. Interstitial macrophage recruitment 24 hours post-CLP was more pronounced in mice undergoing adoptive transfer of memory T cells compared to mice that did not undergo adoptive transfer. IFNγ blockade resulted in higher absolute numbers of T cells, memory T cells, and innate cells in the lungs of Educated mice 24 hours post-CLP suggesting that IFNγ is necessary for curbing an overactive immune response in these mice. In conclusion, the presence of memory T cells affects the course of CLP-induced lung inflammation and may provide a model that more closely resembles sepsis-associated lung injury.

**Summary Statement:** Prior immune memory alters the course of CLP-induced lung inflammation in mice.

## Introduction

Sepsis is defined as life-threatening organ dysfunction caused by a dysregulated host response to infection.^1^ Sepsis is a leading cause of Acute Respiratory Distress Syndrome (ARDS) which is characterized by impaired gas exchange, increased pulmonary interstitial and alveolar fluid, neutrophil recruitment to the lung, loss of type I pulmonary epithelial cells, and hyperplasia of type II pulmonary epithelial cells.^2,3^ Sepsis is the most common extra-pulmonary cause of ARDS and when these two diagnoses occur together, they are associated in increased morbidity and mortality.^4–6^

Cecal ligation and puncture (CLP) is the most widely used murine model of sepsis, but directed therapies that have improved survival following CLP have not been effective in treating human sepsis.^7^ The reasons underlying this failure to translate therapeutic interventions from mice to humans remain unclear. Previous work using CLP has focused on mortality as a primary endpoint, but recent literature suggests that examining organ dysfunction as an outcome variable may improve the translational value of this model.^7^ CLP causes limited pulmonary dysfunction, presenting a clear gap in our ability to model sepsis-associated ARDS.

T cell immune responses have been shown to play a significant role in the development of ARDS in murine models using endotracheal administration of lipopolysaccharide^8^ but the role of T cells in the development of lung injury in CLP is not completely understood. Recent studies suggest that the limited memory T cell repertoire present in laboratory mice may contribute to the inability to translate findings from murine research to humans.^9–12^ We recently demonstrated that inoculation of mice with an anti-CD3ɛ activating antibody induced widespread T cell activation. This process, which we call *Immune Education*, led to the formation of long-term T cell memory in the spleen, liver, and lungs without altering myeloid cell numbers.^13^ *Immune Education* facilitates study of diverse T cell memory responses in isolation (without innate immune changes) in pathogen-free laboratory animals. Previous work demonstrated that *Immune Education* enhances CLP-induced liver dysfunction and mortality. Given the prevalence of lung dysfunction in clinical sepsis, we set out to determine whether *Immune Education* affects the development of pulmonary dysfunction in CLP. In this study, we used *Immune Education* to test the hypothesis that T cell memory alters CLP-induced pulmonary immune responses.

## Methods

### Mice

C57Bl/6J and congenic B6.SJL male mice were obtained from the Jackson Laboratory (Bar Harbor, ME) and were maintained in an AAALAC accredited animal facility at the Feinstein Institutes for Medical Research. Mice were allowed to acclimate to the animal facility for a minimum of 3 days, according to the animal facility guidelines. Wild type C57Bl/6J mice express CD45.2 on all hematopoietic cells while B6.SJL mice express CD45.1. Using both strains in adoptive transfer experiments allowed us to distinguish transferred T cells from native T cells. All studies were approved by the Institutional Animal Care and Use Committee (IACUC #2017-039) and followed National Institutes of Health and Animal Research: Reporting of *In Vivo* Experiments (ARRIVE) guidelines.^14^ To completely assess the effects of CLP on the lung, experiments were performed on several cohorts of mice. Some of the baseline and T24 mice used in these experiments had organs harvested for experiments reported previously,^10,11^ but none of the data included in this report have been previously reported.

### *In vivo* Immunization

We performed *in vivo* T cell activation using Ultra-LEAF anti-mouse CD3ε antibody (50ug, Clone 145-2C11, Biolegend, San Diego, CA) and Ultra-LEAF isotype Armenian Hamster IgG control (50ug, Clone HTK888, Biolegend) as previously described.^13^ Experiments were performed at least 35 days after immunization to allow for the development of T cell memory and to avoid residual acute T cell responses.

### Adoptive Transfer

Adoptive transfer of memory CD4 and CD8 T cells from immunized C57Bl/6J mice to B6.SJL was performed as previously described.^10^ Briefly, we isolated splenic CD4 and CD8 memory (CD90^+^, CD44^+^/CD11a^+^) T cells from Educated C57Bl/6J mice expressing CD45.2 using fluorescence assisted cell sorting. One million CD4 and/or CD8 memory T cells were injected intravenously into congenic Uneducated B6.SJL mice, which express CD45.1. We performed CLP seven days following transfer. Twenty-four hours (hrs.) later, pulmonary cells were isolated and stained for flow cytometry (as described below) to assess leukocyte phenotype.

### Cecal ligation and puncture

We performed double puncture 22-guage CLP in 16-week-old mice as previously described and sacrificed animals at 24, 48 or 72 hrs. post-procedure.^10^ To limit selection bias, the time at which individual mice would be sacrificed was assigned prior to CLP. Mice were fluid resuscitated with subcutaneous normal saline (50 ml/kg) immediately post-operatively and daily for up to 48 hrs. No antibiotics were administered because they can directly impact immune cells^15–18^ and therefore might confound our results. All animals had free access to food and water prior to and after CLP. At the designated time of sacrifice, we euthanized animals with a terminal dose (150mg/kg given intraperitoneally) of pentobarbital sodium and phenytoin sodium.

### *In Vivo* Interferon-gamma blockade

As previously described mice were administered a monoclonal antibody (XMG1.2, 0.5mg, BioXCell, Lebanon, NH) against interferon-gamma (IFNγ) at the time of CLP.^19,20^

### Wet-to-Dry lung weights

Both lungs were harvested following euthanasia. The lungs were weighed immediately, desiccated at 50-56° C, and re-weighed daily for 2-3 days until the weight stabilization indicated full desiccation. Wet-to-dry lung weight ratios were calculated.

### Bronchoalveolar lavage

In a separate pre-designated cohort of mice, we canulated the trachea with a 20-gauge intravenous catheter immediately post-euthanasia. Both lungs were washed three times with the same 1 milliliter (ml) of sterile saline (bronchoalveolar lavage, BAL) and 0.8-1ml of fluid was recovered for collection. Both lungs were then harvested for leukocyte isolation and flow cytometry. BAL protein concentration was quantified using the Pierce BCA Protein assay kit (ThermoFisher, Waltham, MA). The concentrations of 32 cytokines, chemokines, and growth factors in BAL fluid were examined using multiplex analysis (Eve Technologies, Calgary, Alberta).

### Leukocyte Isolation

After BAL, harvested lungs were injected with digestion solution (DNAse, 150 µg/ml, 11284932001 Roche, Sigma-Aldrich, St. Louis, MO; and Type 2 collagenase, 1mg/mL, 17101015, ThermoFisher, Waltham, MA), mechanically disassociated using a scalpel and agitated at 37°C for 45 minutes. Digested tissues were passed through 70 µm filters and cells were washed with phosphate buffered saline. Red blood cells were lysed, and white blood cells were counted using a Countess II Automated Cell Counter (ThermoFisher, Waltham, MA).

### Flow Cytometric Analysis

Following isolation, single cell suspensions were stained for flow cytometric analysis using LIVE/DEAD fixable viability dye (Life Technologies) and antibodies to the following markers: CD90.2, CD44, CD8a, CD4, CD62L, CD44, NK1.1, Ly6C, CD11a, CD11c, CD11b, Ly6G, CD24, SiglecF, CD64, CD45, CD103, MHCII, TNFα, IL2, IL10, IL12, and IFNγ. A second antibody panel that included the following markers was used to evaluate cells from adoptive transfer experiments: CD90.2, CD62L, CD11a, CD4, CD8a, CD44, CD45.2, CD45.1, TNFα, IFNγ, and IL2. A third antibody panel that included the following markers was used for IFNγ-blockade experiments: CD90.2, CD62L, CD11a, CD4, CD8a, CD44, CD5, Nur77, PD1, and CD127. Full antibody details are available in Supplemental Table 1. All flow cytometric analysis was performed on a Becton-Dickinson LSR Fortessa 16-color cell analyzer (BD Bioscience, San Jose, CA) and analyzed using FlowJo software version 10 (BD Bioscience, San Jose, CA). Myeloid cells in the lungs were identified using the gating strategy published by Misharin et al.^21^ Full gating strategy for T cell subset identification is available in Supplemental Figure 1.

### Cytokine Production Assays

Isolated cell suspensions were stimulated for 4 hrs. with phorbol-myristate-acetate (PMA, 50 ng/ml)/ionomycin (Io, 1.5 µmoles/L) in the presence of Brefeldin A (2 µg/ml). Unstimulated cells that served as controls allowed us to quantify background cytokine production.^10^ Intracellular cytokine concentrations were assessed using flow cytometry.

### Histology

#### Tissue fixation and preparation

In a separate cohort of mice, lungs were inflated with 1ml of a mixture containing 4% paraformaldehyde (PFA) and Tissue Plus Optimal Cutting Temperature embedding medium (OCT) mixed 1:1. Lungs were harvested, fixed for 24 hrs. in 4% PFA followed by 24 hrs. of 15% sucrose. They were subsequently placed in 30% sucrose until they sunk. Lungs were embedded in OCT, frozen on dry ice, and stored at -80°C.^22^ Frozen lungs were cut into 10 µm sections using a cryostat.

#### Hematoxylin and eosin staining and Imaging

Lung slides were stained with hematoxylin and eosin (H&E) (Vector Laboratories, Burlingame, CA). Twenty-five non-overlapping high-power fields (40x magnification) from 3 different sections from each mouse were imaged using the Zeiss Axio Observer 7 (Carl Zeiss microscopy, White Plains, NY).

#### Assessment of Lung Injury

Two independent investigators that were blinded to the allocated group used a modified lung injury scoring system (Supplemental Table 2) ^23,24^ to quantify lung injury from images of H&E-stained tissue. Lung injury scores were calculated for each image and then averaged to assign one score per mouse.

#### Immunofluorescent staining

Lung tissue sections underwent immunofluorescent staining for aquaporin-5 (AQP5; rabbit anti-mouse AQP5 (Abcam, Boston, MA) diluted 1:500 in 5% donkey serum (ThermoFisher, Waltham MA)) and surfactant protein C (SPC; rabbit anti-mouse SPC (Abcam, Boston, MA) diluted 1:2000 in 5% donkey serum). Secondary staining was performed with donkey anti-rabbit IgG conjugated to Alexa Fluor 594 (diluted 1:500 in 5% donkey serum). Slides were visualized using the Zeiss LSM880 Airyscan Confocal Microscope. Ten random non-overlapping 20x magnification images from 3 different lung sections were analyzed with Zen 3.3 blue edition (Carl Zeiss Microscopy, White Plains, NY). Fluorescence was quantified using Image J software (Laboratory for Optical and Computational Instrumentation, University of Wisconsin, Madison, WI).^25^ For AQP5-stained sections, the area of positive immunofluorescence was calculated for each image using Image J. The area of immunofluorescence in each image was used to calculate one “average area” of AQP5 fluorescence per mouse. For SPC-stained sections, the number of SPC-stained cells were counted manually for each image. The number of cells per image was averaged to assign one “average number of SPC^+^ cells” per mouse.

### Statistical Analysis

Data were analyzed using 2-way analysis of variance (ANOVA) with Sidak’s Test for Multiple Comparisons (Prism 7.0; GraphPad, San Diego, CA) with a significance level set at p < 0.05. Data are reported as mean ± standard deviation. All data supporting the findings of this study are available within the paper and its supplementary material.

## Results

### *Immune Education* enhances CLP-driven elevations of pulmonary cytokines

We first sought to determine how *Immune Education* altered the effects of CLP on the inflammatory milieu in the alveoli. We therefore measured the concentrations of 32 cytokines and chemokines in post-CLP alveolar lavage fluid obtained from Educated and Uneducated mice. Cytokines in both Educated and Uneducated T0 mice (not subjected to CLP) were low or undetectable. As shown in Fig. 1, CLP did not significantly affect the alveolar levels of the examined cytokines and chemokines in Uneducated mice. However, we found significant elevations in several cytokines associated with T cell immune responses in Educated mice when compared to Uneducated animals post-CLP. Specifically, elevations were noted in IFNγ and IL-17, which are produced by activated T cells; IP-10, GM-CSF and IL-12p70, all IFNγ-responsive cytokines/chemokines; and TNFα, IL1β, and IL-6, which have major roles in promoting immune responses in CLP and human sepsis. Educated mice also had higher concentrations of MIP-1β, which activates respiratory bursts in monocytes and drives T cell responses^26^, and the neutrophil chemokines MCP-1^26^, MIP-2^27^ and KC^28^, which affect innate immune cell function and promote neutrophil recruitment to the lung, a hallmark of ARDS. Interestingly, IL-13, an anti-inflammatory cytokine produced by alveolar macrophages,^29^ was also higher at T24 in Educated mice.

**Figure 1.**
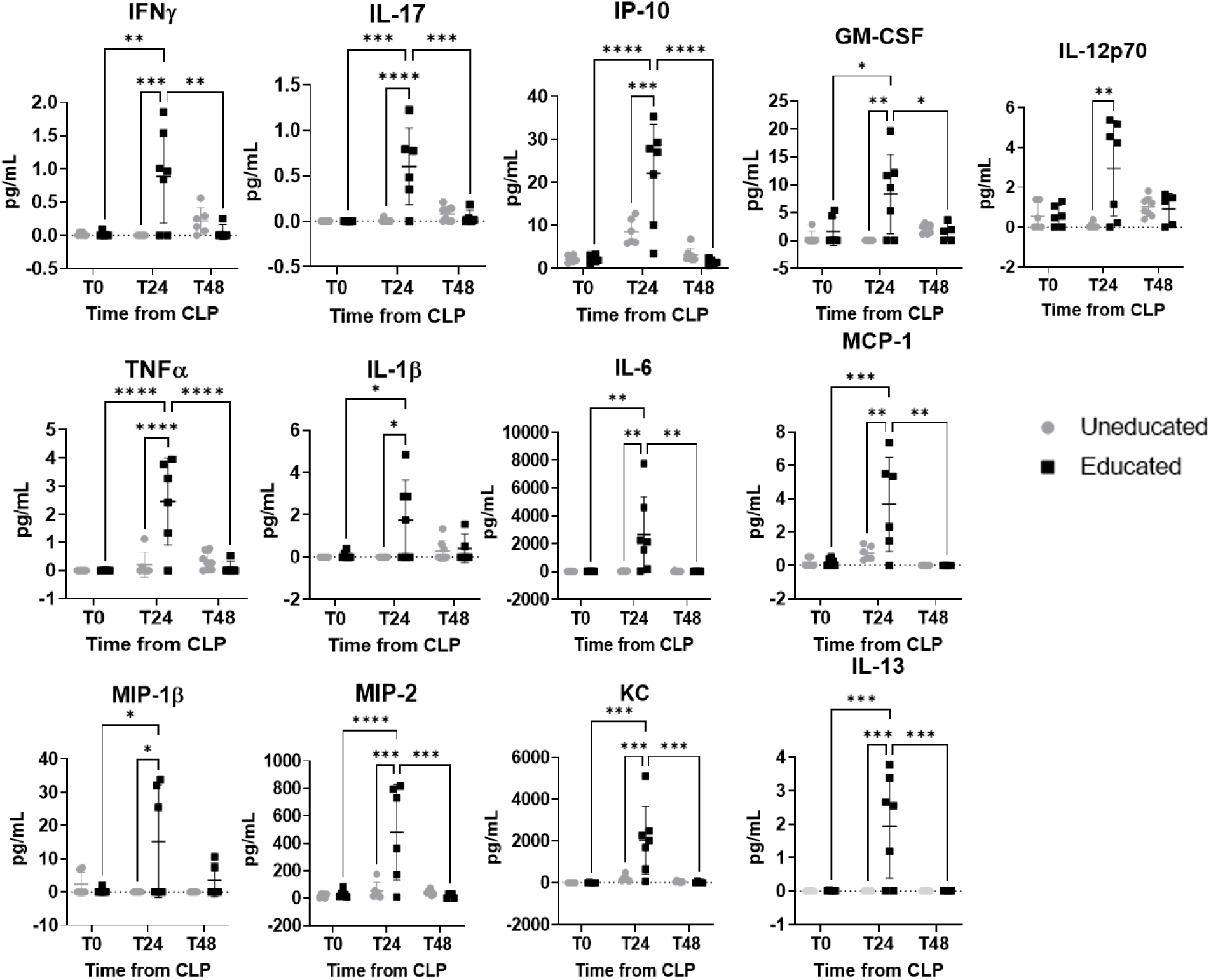
Effects of *Immune Education* on BAL Fluid Protein Levels post-CLP. BAL fluid was collected at the time of euthanasia from Uneducated and Educated mice at T_0_, T_24_, and T_48_ after CLP. BAL cytokine, chemokine, and growth factor concentrations were determined using multiplex ELISA (Eve Technologies). Comparison of BAL protein concentrations at T_0_, T_24_, and T_48_ post CLP in Uneducated (gray circles) and Educated (black squares) mice. n = 5-8 mice per group. Differences in BAL protein concentrations were analyzed by 2-way ANOVA with Sidak’s correction for multiple comparisons. BAL proteins with significant differences between Uneducated and Educated mice are shown. Significance was set at p < 0.05. * p < 0.05; ** p < 0.01; *** p < 0.001, **** p < 0.0001.

### Post-CLP numbers of pulmonary T cells were lower than baseline in both Immune-Educated and Uneducated mice

Educated mice demonstrated an enhanced cytokine response to CLP at the alveolar level (Fig. 1), indicating an enhanced cellular immune response to CLP in association with T cell memory. To assess the direct effects of T cell memory on the pulmonary cellular immune response, we pursued flow cytometric analysis of lungs from Educated and Uneducated animals following CLP. As previously noted, the proportion of memory T cells (CD90^+^, CD11a^+^/CD44^+^) in the lungs at baseline was higher in Educated mice than in Uneducated mice (data not shown).^13^ As previously described in the spleen, baseline numbers of pulmonary CD8 T cells were lower in Immune-Educated mice than in Uneducated animals (Fig. 2A).^13^ Pulmonary T cell numbers were lower at T48 and T72 post-CLP, similar to findings in the spleen,^10,30^ but these differences were similar in both Immune-Educated and Uneducated mice.

**Figure 2.**
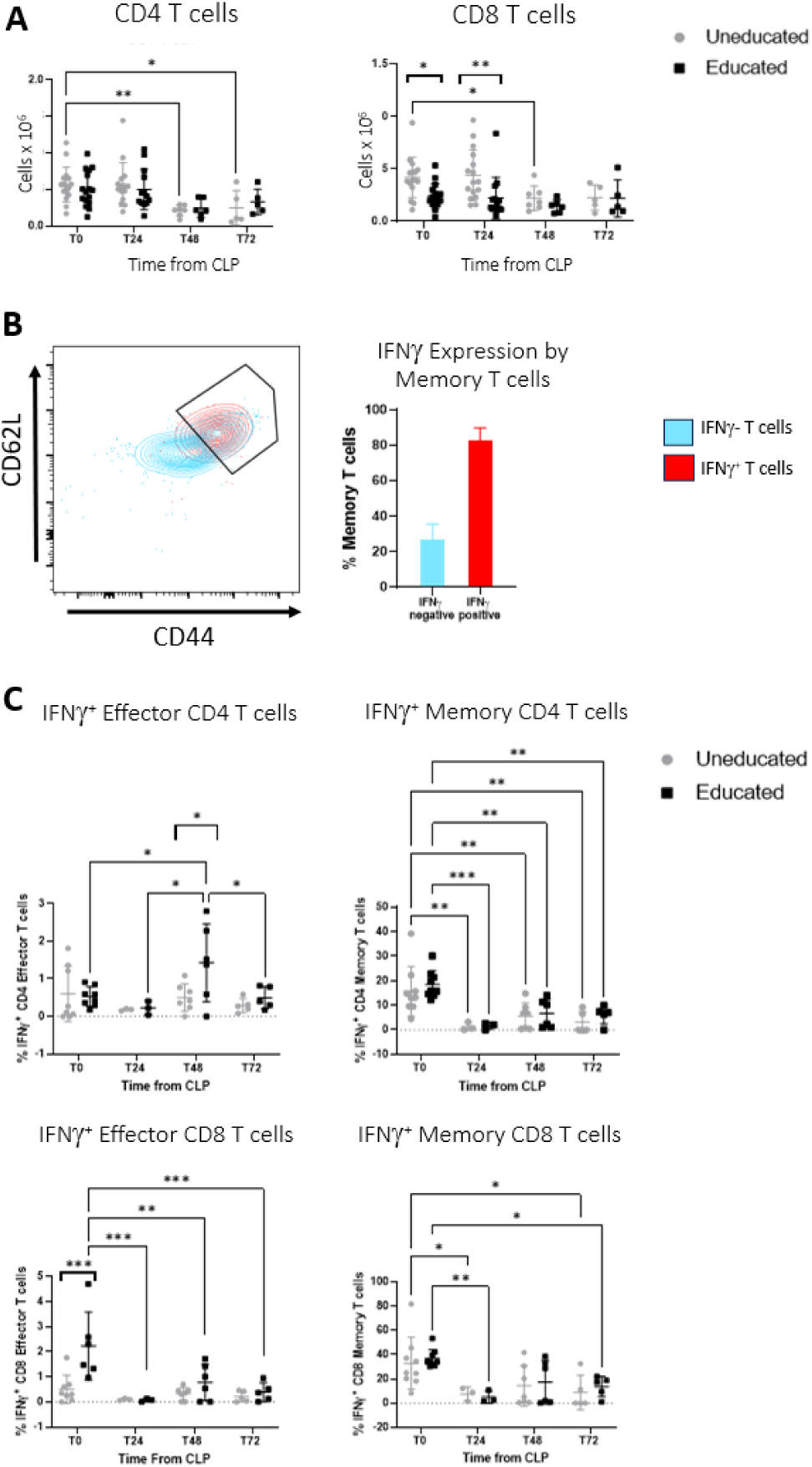
Effects of *Immune Education* on CLP-induced Differences in the Proportion of T cell Subtypes and IFNγ production. Mice were sacrificed at T_0_, T_24_, T_48_ and T_72_ after CLP. Lungs were harvested, digested, and single cells were stained for flow cytometry. **A.** The graphs show absolute number of Lung CD4 T cells (CD90^+^, CD4^+^) and CD8 T cells (CD90^+^, CD8^+^) in Uneducated (gray circles) and Educated (black squares) mice after CLP. n = 5-16 mice per group. **B and C.** Pulmonary T cells were stimulated *ex vivo* with phorbol-myristate-acetate/ionomycin in the presence of brefeldin A and cytokine production was quantified using flow cytometry. **B.** Contour plot showing IFNγ^+^ T cells (red) overlying memory T cells (CD44^+^/CD62L^+^) in a representative sample of an Immune-Educated mouse at T_0_. IFNγ^+^ cells were predominantly memory T cells. The bar graph shows the proportion of memory T cells producing IFNγ (in red) compared to the proportion of memory T cells not producing IFNγ (in blue). **C.** Proportions of IFNγ-producing lung CD4 and CD8 Memory (CD44^+^/CD11a^+^) and Effector (CD44^-^/CD11a^-^) T cells in Uneducated (gray circles) and Educated (black squares) mice after CLP. n=3-8 mice per group. Significance was determined using 2-way ANOVA with Sidak’s correction for multiple comparisons and was set at p < 0.05. * p < 0.05; ** p < 0.01; *** p < 0.001.

### *Immune Education* augmented IFNγ^+^ effector CD4 T cell responses to CLP

Despite lower pulmonary T cell numbers, the cytokine response observed in alveolar fluid was significantly augmented in Educated mice following CLP (Fig. 1), suggesting that functional responses to CLP may be augmented by T cell memory. IFNγ, produced by T cells and NK cells, links adaptive and innate cell responses through macrophage and neutrophil stimulation, leukocyte attraction, enhancement of NK cell activity, and regulation of B cell functions.^31,32^ Post-CLP alveolar fluid levels of the IFNγ-responsive proteins IP-10, GM-CSF and IL-12p70 were higher in Educated than in Uneducated animals, suggesting that *Immune Education* enhanced post-CLP IFNγ activity. We therefore compared the effects of *Immune Education* on CLP-induced expression of IFNγ in response to *ex vivo* stimulation with PMA/Io. Results are detailed in Fig. 2B and 2C. IFNγ was primarily produced by pulmonary memory T cells (CD90^+^, CD44^+^/CD62L^+^) (Fig. 2B). Prior to CLP, Immune-Education did not alter the percentage of CD4^+^ effector (CD90^+^, CD4^+^, CD44^-^/CD11a^-^) or memory T cells (CD90^+^, CD4^+^, CD44^+^/CD11a^+^) that expressed IFNγ (Fig. 2C) when compared to Uneducated controls. The percentage of IFNγ^+^ memory CD4 T cells was lower than baseline at all post-CLP time points in both Educated and Uneducated animals. However, 48 hours post-CLP, the proportion of IFNγ^+^ effector CD4 T cells in Immune-Educated mice was higher than at baseline and compared to Uneducated mice, indicating a T effector response with significant activation at this timepoint post-CLP in Educated mice that is absent in Uneducated mice.

When CD8 T cells were examined, further changes in the T cell IFNγ response were noted. Again, no differences in the percentage of IFNγ^+^ memory CD8 T cells between Educated and Uneducated animals could be detected at any specific time point studied. *Immune Education* did induce differences between Educated and Uneducated mice in the *ex vivo* expression of IFNγ by effector CD8 T cells at baseline. Prior to CLP, the proportion of IFNγ^+^ effector CD8 T cells in the lung were higher in Educated than in Uneducated mice, indicating an enhanced potential IFNγ response at the time of CLP. The proportion of IFNγ^+^ effector CD8 T cells was lower post-CLP in Educated mice whereas the CD8 T cell IFNγ response was absent in Uneducated animals both at baseline and following CLP (Fig 2B).

### *Immune Education* augmented early inflammatory and late reparative pulmonary innate immune responses to CLP

IFNγ drives monocyte, macrophage, and neutrophil responses during infection and inflammation.^31,32^ The presence of IFNγ^+^ effector CD8 T cells prior to CLP and the development of IFNγ^+^ effector CD4 T cells by 48 hrs. following CLP in Educated animals suggests that *Immune Education* may license memory T cells to promote Effector T cells to produce IFNγ. To assess effects of T cell memory on CLP-associated innate immune responses in the lung, we examined myeloid cell numbers in Immune-Educated and Uneducated mice (Fig. 3). *Immune Education* did not affect the absolute number of pulmonary myeloid cells at baseline. We found significant differences in macrophage and monocyte numbers when Educated and Uneducated mice were compared. Interstitial macrophages are pro-inflammatory and promote neutrophil recruitment whereas alveolar macrophages are involved in lung repair and can limit neutrophil infiltration.^21^ At 24 hrs. post-CLP, the number of interstitial macrophages (CD45^+^, CD11b^hi^, MHCII^+^, CD64^+^) was higher than T0 in both cohorts, but the difference was more pronounced in Educated mice. Differences from T0 in interstitial macrophage numbers were no longer present in either cohort at later post-CLP timepoints (Fig. 3). Ly6C^+^ inflammatory monocyte (CD45^+^, CD11b^hi^, MHCII^-^, Ly6C^+^) numbers were also higher at T24 than T0 in Immune-Educated mice – no change from T0 was noted in Uneducated mice. By 48 hrs. post-CLP, the numbers of Ly6C^+^ inflammatory monocytes were higher than T0 in both cohorts of mice and remained higher at 72 hrs. *Immune Education* did not significantly affect these post-CLP differences (Fig. 3). In contrast, the absolute numbers of alveolar macrophages (CD45^+^, SiglecF^hi^, CD11c^+^/CD64^+^) in Educated mice 72 hrs. post-CLP were higher than at T0 and were significantly different from the numbers in Uneducated animals (Fig. 3). Alveolar macrophage numbers did not differ from T0 at any time point after CLP in Uneducated mice; there were also no differences at T24 and T48 in Educated mice. Given post-CLP differences in pro- and anti-inflammatory myeloid cell numbers, we calculated the ratio of Ly6C^+^ inflammatory monocytes to alveolar macrophages to quantify the inflammatory to anti-inflammatory balance.^33^ This number was higher at 24 and 48 hrs. post-CLP in Educated, but not Uneducated, mice. Seventy-two hrs. after CLP the pattern was reversed; the ratio was significantly higher than T0 in Uneducated but not Educated mice. Taken together, these data suggest that *Immune Education* is associated with enhanced inflammation in the lungs 24 and 48 hrs. post-CLP. By seventy-two hrs. post-CLP, this pattern is reversed towards reparative immune responses in Educated mice, while Uneducated mice are only beginning to demonstrate inflammation after 48 hrs.

**Figure 3.**
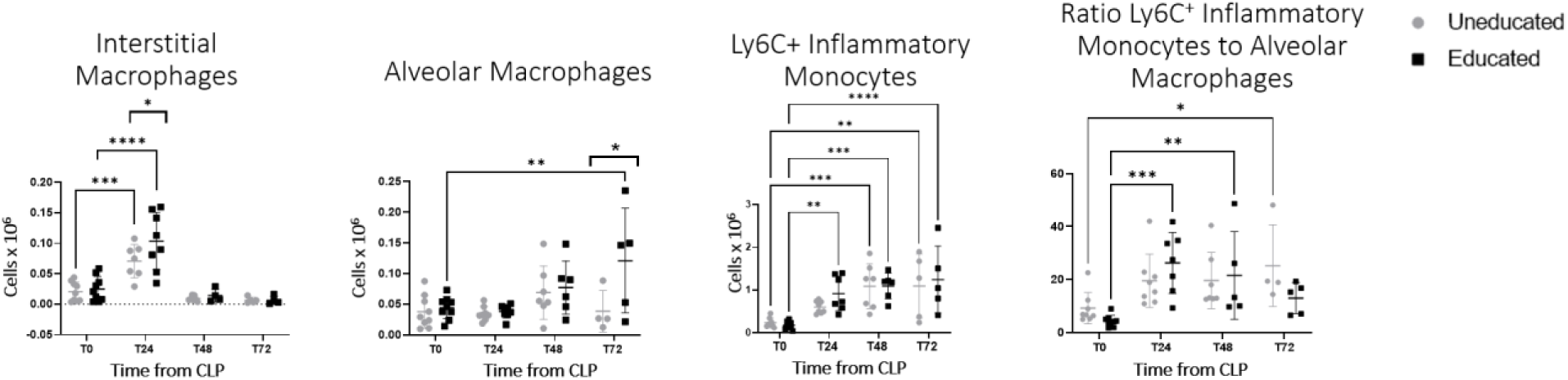
Effects of Immune Education on CLP-induced Differences in Pulmonary Monocytes and Macrophages. Mice were sacrificed at T_0_, T_24_,T_48_, and T_72_ after CLP. Lungs were harvested, digested, and single cells were stained for flow cytometry. Absolute numbers of lung interstitial macrophages (CD45^+^, CD11b^hi^, MHCII^+^, CD64^+^), alveolar macrophages (CD45^+^, SiglecF^hi^, CD11c^+^/CD64^+^), and Ly6C^+^ inflammatory monocytes (CD45^+^, CD11b^hi^, MHCII^-^, Ly6C^+^) in Uneducated (gray circles) and Educated (black squares) mice after CLP. Ratio of Ly6C^+^ inflammatory monocytes to alveolar macrophages in Uneducated and Educated mice after CLP is shown. n=4-8 mice per group. Significance was determined using 2-way ANOVA with Sidak’s correction for multiple comparisons and was set at p < 0.05. * p < 0.05; ** p < 0.01; *** p < 0.001; **** p < 0.0001.

### Intrinsic T cell memory functions drive higher pulmonary recruitment of interstitial inflammatory macrophages

To test whether augmented innate immune responses are due to the presence of T cell memory, adoptive transfer of splenic CD4 memory T cells (CD4^+^, CD11a^+^/CD44^+^), splenic CD8 memory T cells (CD8^+^, CD11a^+^/CD44^+^), or both from Immune-Educated mice to B6.SJL mice was used. Five groups of mice were studied: 1) B6.SJL mice not subjected to CLP or to adoptive transfer (untreated T0), 2) B6.SJL mice subjected to CLP but not to adoptive transfer (untreated T24), 3) B6.SJL mice treated with CD4 memory T cells and subjected to CLP (CD4 treated T24), 4) B6.SJL mice treated with CD8 memory T cells and subjected to CLP (CD8 treated T24), and 5) B6.SJL mice treated with CD4 and CD8 memory T cells and subjected to CLP (CD4 + CD8 treated T24). Like data obtained from Immune-Educated mice (Fig. 3), at 24 hrs. post-CLP, mice treated with adoptive transfer of either memory CD4 or memory CD8 T cells had a higher percentage of interstitial macrophages than untreated mice at T0 and untreated mice 24 hrs. post-CLP (Fig. 4), suggesting that intrinsic T cell memory functions drive pulmonary recruitment of interstitial macrophages. Interestingly, this difference was not significant in animals receiving both CD4 and CD8 memory T cells. At 24 hrs. post-CLP, the proportions of Ly6C^+^ inflammatory monocytes and alveolar macrophages in animals pre-treated with any combination of splenic CD4 and CD8 memory T cells were not significantly different from proportions in untreated mice.

**Figure 4.**
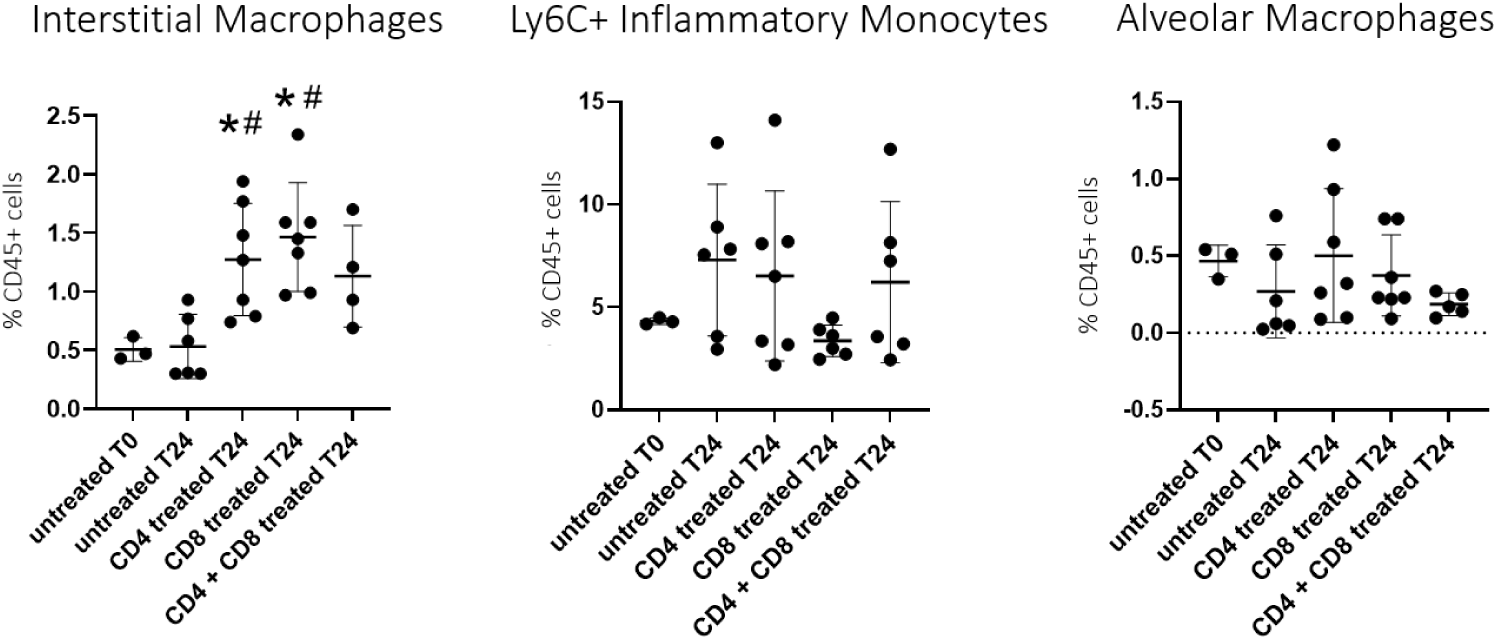
Effects of *Immune Education* through Adoptive Transfer of Memory T cells on CLP-induced differences in Pulmonary Myeloid Cell numbers. Effects of CLP on the proportions of lung interstitial macrophages (CD45^+^, CD11b^hi^, MHCII^+^, CD64^+^), Ly6C^+^ inflammatory monocytes (CD45^+^, CD11b^hi^, MHCII^-^, Ly6C^+^), and alveolar macrophages (CD45^+^, SiglecF^hi^, CD11c^+^/CD64^+^) in B6.SJL mice who underwent adoptive transfer of memory CD4 T cells, memory CD8 T cells, or both from Immune-Educated C57Bl/6 mice compared to no CLP and untreated T_24_. n=3-7 mice per group. Significant was determined using 1- way ANOVA with Sidak’s correction for multiple comparisons and was set at p < 0.05, * p < 0.05 compared to untreated T_0_; # p < 0.05 compared to untreated T_24_.

### *Immune Education* accelerates the development of CLP-induced lung injury in mice

Considering the enhanced inflammatory immunologic response observed with *Immune Education* following CLP (Fig. 1-3), we hypothesized that lungs from Immune Educated animals following CLP would demonstrate aspects of enhanced injury. Qualitative histological differences between Educated and Uneducated mice following CLP were evident on representative H&E-stained sections (Fig. 5A). When compared to Uneducated animals, Educated mice had more neutrophilic infiltration, a greater accumulation of proteinaceous debris in the alveolar space, and alveolar septal thickening. After CLP, *Immune Education* significantly enhanced overall lung injury scores. Specifically, lung injury scores peaked in Educated mice 48 hrs. after CLP when compared to T0 (Fig. 5B). Seventy-two hrs. after CLP, lung injury scores were higher than T0 in both Educated and Uneducated mice. Expression of aquaporin 5 (AQP5, a marker used to quantify the number of type 1 pulmonary epithelial cells^7,34^) in Educated mice was not different from T0 at all post-CLP time points, while in Uneducated animals, AQP5 was significantly lower than T0 at 24 and 48 hrs. post-CLP. There were no differences in AQP5 expression between Educated and Uneducated mice at any time point. At T24, numbers of cells containing surfactant protein C (SPC, expressed in type II pneumocytes^7^), were significantly higher than T0 in both Educated and Uneducated mice; no difference from T0 was noted at T48 in either group (Fig. 5C). No CLP-induced differences in wet-to-dry lung weights or in total BAL protein concentration (data not shown) were noted; these findings suggest that pulmonary edema was minimal. Even though specific markers of lung injury were not enhanced by *Immune Education*, these findings demonstrate an overall enhancement of lung inflammation in Immune Educated CLP.

**Figure 5.**
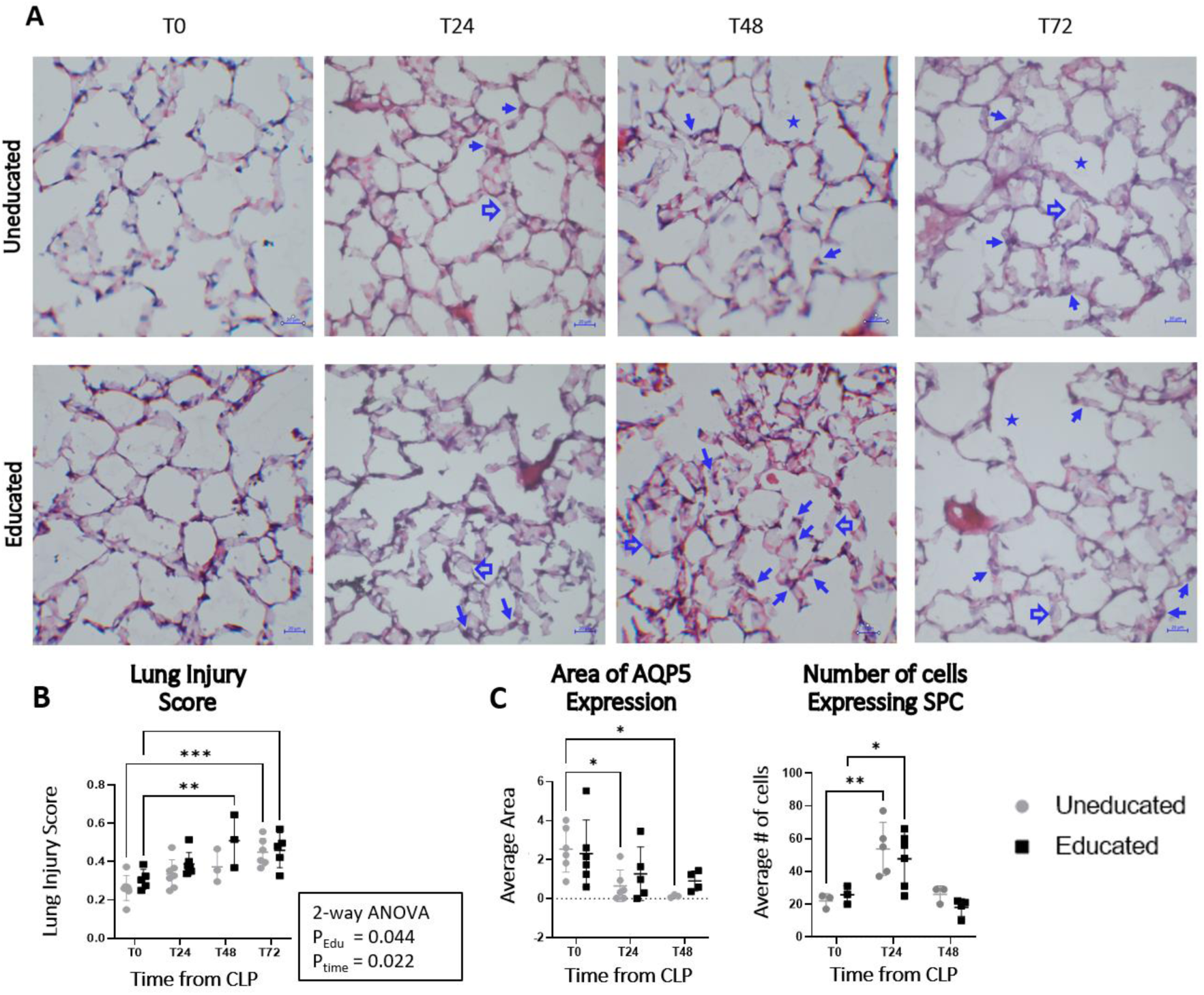
Effects of CLP on Lung Injury in Educated and Uneducated Mice. **A.** Representative images from H&E stains of Uneducated and Educated mice at T_0_, T_24_, T_48_, and T_72_ taken at 40x magnification. Closed arrows indicate neutrophils, star indicates proteinaceous debris, and open arrows indicate alveolar septal thickening. **B.** Lung injury scores were calculated from H&E-stained samples as shown in supplemental table 2. Data represent individual lung injury scores/mouse. Scores determined by averaging results from 25 non-overlapping 40X magnification area/fixed sections, 3 fixed sections/mouse. Comparison of lung injury scores at T_0_, T_24_, T_48_, and T_72_ post CLP in Uneducated (gray circles) and Educated (black squares) mice. n = 3-7 mice in each group. p-values indicate results from 2-way ANOVA showing effects *of Immune Education* (P_Edu_) and of time from CLP (P_time_) **C.** Aquaporin 5 (AQP5) and Surfactant Protein C (SPC) abundance at T_0_, T_24_ and T_48_ post-CLP in Uneducated (grey circles) and Educated (black squares) mice. Staining as detailed in the methods. AQP5 - Data represent individual values of positive immunofluorescence/mouse. Area determined by averaging results after examining 10 non-contiguous 20X magnification areas/fixed section, 3 fixed sections/mouse. Quantification using image J software (Laboratory for Optical and Computational Instrumentation (LOCI), Madison, WI). SPC - Data represent individual values of manually counted SPC-stained cells/mouse. Numbers determined by averaging results after examining 10 non-contiguous 20X magnification areas/fixed section, 3 fixed sections/mouse. Quantification using image J software (LOCI, Madison, WI). Mean - horizontal bar, standard deviation – error bars. n = 3-6 mice per group. Significance was determined using 2-way ANOVA with Sidak’s correction for multiple comparisons and was set at p < 0.05. * p < 0.05; ** p < 0.01; *** p < 0.001.

### Interferon-gamma blockade enhances CLP-induced increases in T cell numbers and T cell activation in Educated mice

Lung inflammation in Educated mice was preceded by higher induced IFNγ production by CD8 effector T cells at T0 and higher proportions of lung effector IFNγ^+^ CD4 T cells following CLP. These results reflect higher alveolar fluid levels of IFNγ and IFNγ-inducible cytokine levels in Educated mice 24 hrs. following CLP. We hypothesized that in Immune-Educated mice, early increased T cell IFNγ production drives early myeloid cell recruitment to lung tissue. To test this hypothesis, we blocked IFNγ in Immune-Educated and Uneducated mice at the time of CLP and assessed immune responses 24 hrs. later. We found that, compared to Uneducated mice and Educated mice that did not undergo IFNγ blockade, Educated mice that underwent IFNγ blockade had higher numbers of total T cells, central memory T cells (CM, CD90^+^, CD44^+^/CD62L^+^) and effector memory T cells (EM, CD90^+^, CD44^+^/CD62L^-^). Both CD4 and CD8 T cells numbers were higher. IFNγ blockade led to significant enhancement of markers of T cell activation including Nur77, CD127, CD5, and PD1 (Fig. 6), but had no effect on CD69 expression. Nur77 expression is an early marker of T cell receptor activation.^35,36^ CD127, or interleukin-7 receptor α (IL-7Rα), is important for promoting T cell maturation and survival of memory T cells.^37–39^ CD5^+^ T cells have a higher affinity for self-antigens^40,41^ and are often elevated in the setting of increased IL-7 responsiveness^42^ which is consistent with our finding of higher CD127^+^ T cells. Programmed cell death protein 1 (PD1) is another early activation marker and an immune checkpoint regulator that attempts to prevent excessive T cell activation, allowing for the maintenance of tolerance and preventing autoimmunity. It is typically elevated when the host experiences significant antigen exposure and plays a major role in promoting T cell hypo-reactivity when antigen is present for long periods such as during chronic infection or autoimmune disease.^43^ The ratio of CD5:PD1 was calculated to assess whether there was a skew towards or away from a risk of autoreactivity. This ratio was significantly lower in Educated mice at T24, but it was not affected by IFNγ blockade.

**Figure 6.**
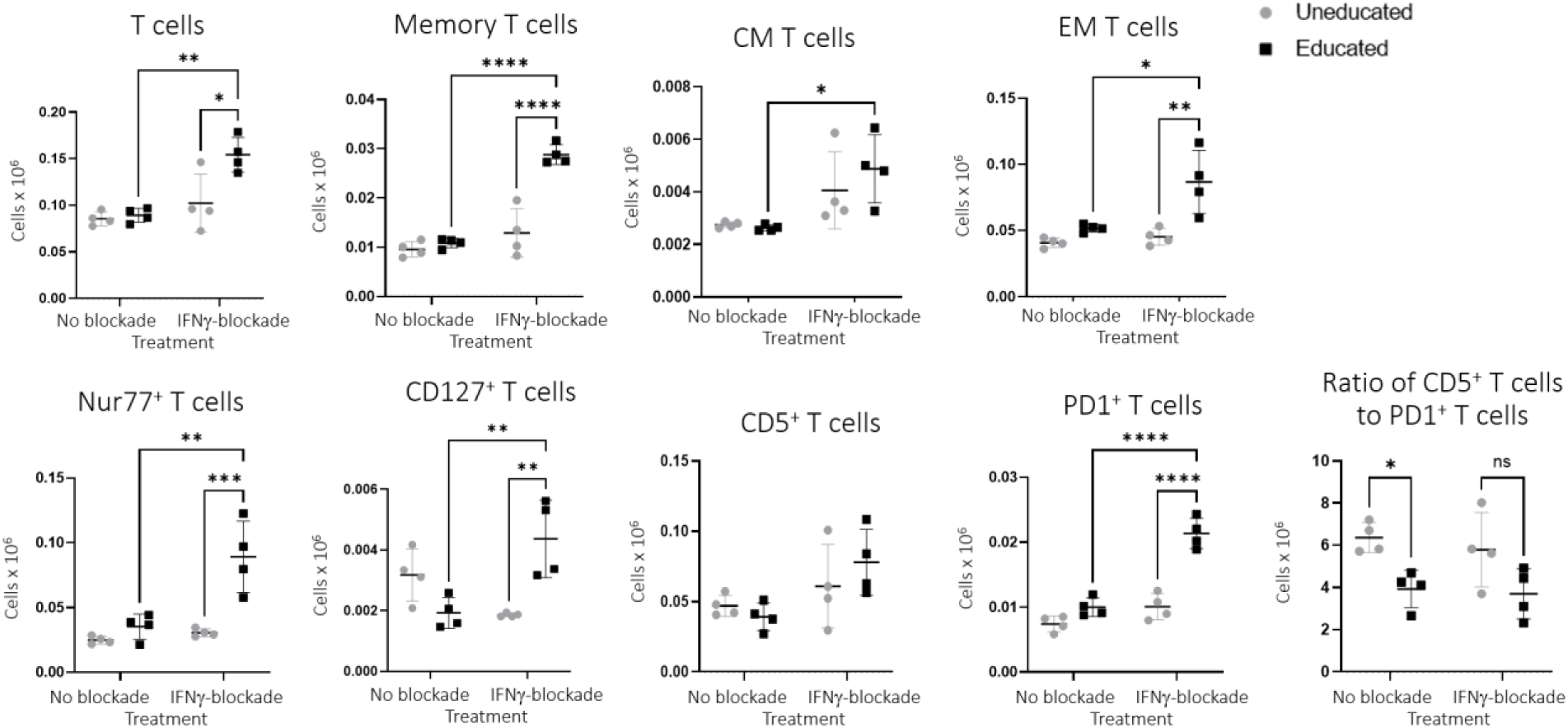
Effects of IFNγ blockade on CLP-induced pulmonary T cell numbers in Immune-Educated and Uneducated mice. Mice underwent CLP and simultaneous IFNγ blockade or vehicle control. Mice were sacrificed at T_24_ after CLP. Lungs were harvested, digested, and single cells were stained for flow cytometry. The graphs show absolute number of Lung T cells (CD90^+^), memory T cells (CD90^+^, CD44^+^/CD11a^+^), central memory T cells (CM, CD90^+^, CD44^+^/CD62L^+^), effector memory T cells (EM, CD90^+^, CD44^+^/CD62L^-^), Nur77^+^ T cells (CD90^+^, Nur77^+^), CD127^+^ T cells (CD90^+^, CD127^+^), PD1^+^ T cells (CD90^+^, PD1^+^), and CD5^+^ T cells (CD90^+^, CD5^+^) in Uneducated (gray circles) and Educated (black squares) mice. n = 4-5 mice per group. Significance determined using 2-way ANOVA with Sidak’s correction for multiple comparisons and was set at p < 0.05. * p < 0.05; ** p < 0.01, *** p < 0.001, **** p < 0.0001.

### Interferon-gamma blockade results in higher CLP-induced recruitment of pulmonary innate cells in Educated Mice

IFNγ prevents upregulation of markers of T cell activation, and specifically of T cell receptor activation, following CLP. T cell receptor activation occurs in response to interactions with antigen presenting cells including macrophages, monocytes and dendritic cells. These findings may indicate that IFNγ blockade in CLP allows enhanced innate immune responses due to lack of suppression by IFNγ, and leads to higher T cell activation due to interactions with antigen presenting cells. We assessed whether the CLP-induced pulmonary T cell changes noted in Educated mice after IFNγ block were associated with changes in pulmonary myeloid cells. Compared to Uneducated mice and Educated mice that did not undergo IFNγ block, Educated mice who underwent IFNγ blockade had higher absolute numbers of the following cell types: alveolar macrophages, interstitial macrophages, Ly6C^+^ inflammatory monocytes, Ly6C^-^ patrolling monocytes (CD45^+^, CD11b^hi^, MHCII^-^, Ly6C^-^), neutrophils (CD45^+^, CD11b^+^/Ly6G^+^), eosinophils (CD45^+^, Ly6G^-^, CD11c^+^/SiglecF^+^), CD11b^+^ dendritic cells (CD45^+^, CD11b^hi^, MHCII^+^, CD64^-^/CD24^+^), CD103^+^ dendritic cells (CD45^+^, CD11c^+^/CD24^+^) (Fig. 7). These results suggest that in Educated mice, T cell IFNγ is necessary to stem innate immune responses that are otherwise unchecked without this cytokine but may simultaneously contribute to an overall pulmonary inflammatory response to CLP.

**Figure 7.**
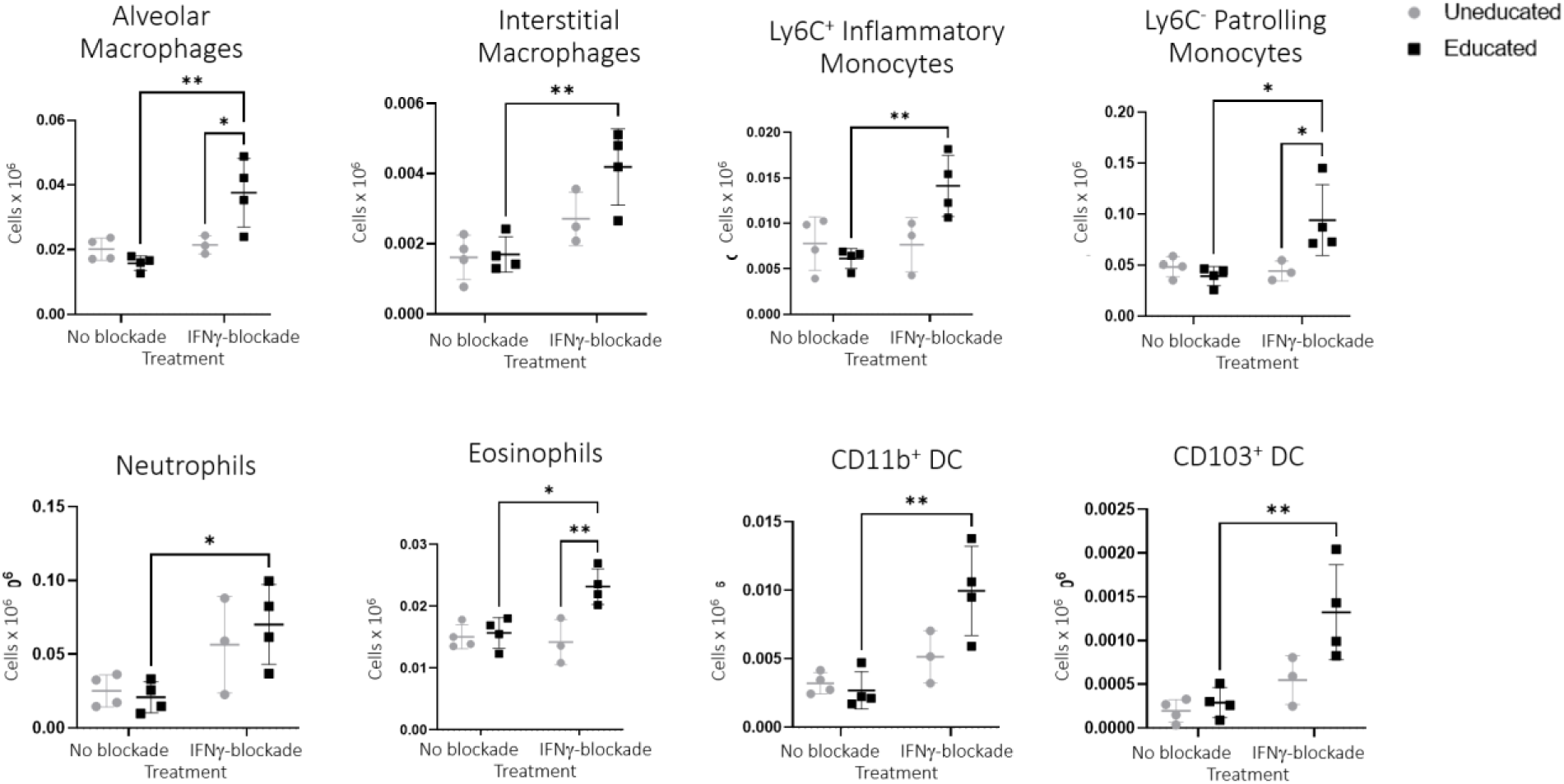
Effects of IFNγ blockade on CLP-induced pulmonary myeloid cell numbers in Immune-Educated and Uneducated mice. Mice underwent CLP and simultaneous IFNγ blockade or vehicle control. Mice were sacrificed at T_24_ after CLP. Lungs were harvested, digested, and single cells were stained for flow cytometry. The graphs show absolute number of alveolar macrophages (CD45^+^, SiglecF^hi^, CD11c^+^/CD64^+^), interstitial macrophages (CD45^+^, CD11b^hi^, MHCII^+^, CD64^+^), Ly6C^+^ inflammatory monocytes (CD45^+^, CD11b^hi^, MHCII^-^, Ly6C^+^), Ly6C^-^ patrolling monocytes (CD45^+^, CD11b^hi^, MHCII^-^, Ly6C^-^), neutrophils (CD45^+^, CD11b^+^/Ly6G^+^), eosinophils (CD45^+^, Ly6G^-^, CD11c^+^/SiglecF^+^), CD11b^+^ dendritic cells (CD45^+^, CD11b^hi^, MHCII^+^, CD64^-^/CD24^+^), CD103^+^ dendritic cells (CD45^+^, CD11c^+^/CD24^+^) in Uneducated (gray circles) and Educated (black squares) mice. n = 4-5 mice per group. Significance determined using 2-way ANOVA with Sidak’s correction for multiple comparisons and was set at p < 0.05. * p < 0.05; ** p < 0.01.

## Discussion

The data presented here indicate that *Immune Education* augments the pulmonary inflammatory response to CLP. At baseline *Immune Education* stimulated lung memory T cell development, with a greater proportion of cells being lung effector IFNγ^+^ CD8 T cells despite fewer total CD8 T cells. These effects were associated with a CLP-induced inflammatory response that differed from what developed in Uneducated animals. Specifically, the abundance of alveolar inflammatory cytokines and chemokines (with a strong IFNγ signature), the number of interstitial macrophages, and the ratio of Ly6C^+^ inflammatory monocytes to alveolar macrophages were significantly higher at T24 than at T0 in Educated mice, and significantly higher than values in Uneducated mice. Increased numbers of myeloid cells, specifically interstitial macrophages, can be recapitulated in adoptive transfer experiments indicating that the data seen are immune responses directly evoked by memory T cells. Importantly, alveolar cytokine concentrations indicate that IFNγ contributed substantially to the differences. Overall, these results show that T cell memory augments the pulmonary immune response to CLP after 24 hrs.

IFNγ blockade experiments suggest that IFNγ prevents further spreading of the inflammatory response to CLP in the lungs of Educated mice. IFNγ blockade also resulted in an expansion of memory T cells in Educated mice which may have been part of a compensatory mechanism attempting to increase IFNγ secretion. Conversely, IFNγ is essential in inducing cytolytic activity when T cells interact with antigen presenting cells. Blockade of IFNγ may have prevented antigen presenting cell pruning by CD4 and CD8 T cells, allowing spreading of T cell activation.^44,45^ In fact, we found higher numbers of T cells expressing markers of T cell activation (Nur77, CD127, and CD5) and PD1 after IFNγ blockade. These findings are consistent with the paradigm that IFNγ prevents the spreading of T cell activation. Despite increased CLP-induced T cell activation following IFNγ blockade, there was no effect on CD69 expression. CD69 expression can be triggered by diverse stimuli and is not limited to activation via the T cell receptor.^46,47^ IFNγ blockade did enhance Nur77 expression, which requires T cell receptor activation, suggesting that IFNγ prevents T cell receptor activation in the lungs and decreases cell-cell interactions. CD5 and PD1 expression were examined because of emerging evidence suggesting that sepsis can induce autoimmune-like processes through the release of damage-associated molecular patterns from injured tissues that trigger immune responses against self-antigens, and through the dysregulation of immune checkpoints.^48–51^ CD5 expression increases the risk of T cell activation by autoantigens and PD1 expression prevents T cell activation. The ratio of CD5:PD1 was significantly lower in Educated mice, but it was not affected by IFNγ blockade suggesting that there is lower risk for autoreactivity in Educated mice exposed to CLP that is independent from the action of IFNγ. Lower risk of autoreactivity could contribute to curbing the dysregulated inflammatory response in the lung and may explain why there are signs of earlier recovery from lung injury in Educated mice.

Forty-eight hrs. after CLP, we noted a greater proportion of lung effector IFNγ^+^ CD4 T cells in Educated mice compared to Uneducated. This finding along with the higher concentration of pro-inflammatory alveolar proteins in Educated mice 24 hrs. after CLP suggests that *Immune Education* accelerated the development of pulmonary inflammation. Immune-Educated mice also had higher levels of the immune-regulatory proteins IL-13^29^ and GM-CSF^52,53^ in their alveolar fluid 24 hrs. post CLP. Elevations in these cytokines may indicate a counter-regulatory response in Educated mice, limiting the level of lung inflammation. IFNγ blockade experiments suggest that IFNγ elevations at earlier timepoints in Educated mice, may also be curbing the inflammatory response to CLP. This interpretation is supported by the higher number of alveolar macrophages present in Educated mice 72 hrs. post-CLP. The prominent phenotype of alveolar macrophages favors tissue repair rather than inflammation^33^. We speculate that higher alveolar macrophage numbers at T72 may reflect the increased alveolar GM-CSF concentrations noted two days earlier^54^. In sum, inflammation occurs early post-CLP in Educated mice, and *Immune Education* appears to facilitate recovery after 72 hrs.

Despite differences in lung inflammation and lung injury scores after CLP with *Immune Education*, AQP5 and SPC staining were similar in Educated and Uneducated mice. AQP5 staining, a marker of alveolar type I cells^34^, is lower than baseline early in the course of ARDS^55^. We identified less AQP5 staining post-CLP than at baseline in Uneducated mice but did not detect a similar difference in Educated mice. Further, a loss of type I cells is often followed by proliferation of type II pneumocytes, usually identified by increased levels of SPC. If this response is limited to a monolayer, it is likely part of an adaptive repair mechanism^55^. However, the presence of multiple layers of type II cells is usually mal-adaptive and associated with fibrosis. Expression of SPC was higher than baseline at T24 in both Immune-Educated and Uneducated mice, but whether this proliferation was adaptive or pathologic was not examined and should be investigated in the future. The disconnection of these markers from the overall inflammatory score indicates that different mechanistic processes may drive each of these components of pulmonary inflammatory injury.

Sepsis is the most common cause of ARDS^4–6^, yet pulmonary dysfunction following CLP does not fully recapitulate human ARDS^7^. Recent evidence indicates that laboratory mice lack much of the well-developed T cell memory present in adult humans^9^. In our previous work, we demonstrated that inducing *Immune Education* (that is, activating CD4 and CD8 T cells and producing long-lasting T cell memory) via treatment of mice with anti-CD3ɛ activating antibody^13^ enhanced CLP-induced inflammatory responses, liver injury, and mortality^10^. The experiments described here suggest that *Immune Education* also altered CLP-induced inflammation in the lungs, adding to our contention that T cell memory responses are important contributors to development of organ dysfunction that is the sine qua non of sepsis. It is intriguing to speculate that *Immune Education* contributes to the pathogenesis of the recently described “hyperinflammatory sub phenotype” of ARDS^56^.

There are several limitations to this study. First, despite its widespread use, CLP differs significantly from human sepsis^7^. Further, serial post-CLP measurements in individual animals are not possible. Therefore, we can only identify differences between mean values of measured variables at different post-CLP timepoints. Additionally, anti-CD3ε activating antibody induces a non-specific T cell memory that likely does not recapitulate the full complexity of natural immune development^10^, hence it may not reflect the impact of naturally developed T cell memory on disease in mice and may not target the bacteria causing infection in the CLP model. Future studies in naturalized mice may be warranted, though the complexity of changes in the immune system of these animals limits mechanistic studies focused on the T cell response specifically, justifying use of our complimentary current model.

In conclusion, our previous work demonstrated that *Immune Education* alters the murine response to CLP in a manner that affects organ dysfunction and mortality. This study shows that *Immune Education* also alters the pulmonary immune response to CLP. These findings highlight the importance of immune memory in using laboratory animal models to recapitulate human immune responses to infectious stimuli and add to our evolving understanding of the contribution of T cell memory to the pathobiology of sepsis.

## Authorship Contribution Statement

MRB, CSD, and MDT substantially contributed to the conceptualization, investigation, data curation, data analysis, project administration, project oversight, and writing of the manuscript.

## Acknowledgements

We would like to thank Li Lou, Mei Qi, Mabel N. Abraham, Omar Yaipen, and Ana Nedeljkovic-Kurepa. LL provided guidance with the methodology and analysis of lung histology. LL and MQ served as blind investigators to score H&E sections using the modified lung injury score. OY assisted in lung tissue fixation and preparation for staining. MA and ANK performed CLP surgeries.

---

**Supplemental Table 1.** Full flow cytometry antibody details.

| Target | Clone | Cat. Number | Company | Fluorophore | Host Species | Target Species |
| --- | --- | --- | --- | --- | --- | --- |
| CD4 | RM4-5 | 100552 | Biolegend | BV785 | Rat | Mouse |
| CD5 | 53-7.3 | 100634 | Biolegend | APC-Cy7 | Rat | Mouse |
| CD8a | 53-6.7 | 100730 | Biolegend | AF700 | Rat | Mouse |
| CD11a | M17/4 | 740676 | BD Bioscience | BV711 | Rat | Mouse |
| CD11b | M1/70 | 101256 | Biolegend | PE/Dazzle594 | Rat | Mouse/Human |
| CD11c | HL3 | 564079 | BD Bioscience | BV650 | Hamster | Mouse |
| CD24 | M1/69 | 101805 | Biolegend | FITC | Rat | Mouse |
| CD44 | IM7 | 61-0441-82 | Invitrogen | PE-eFluor610 | Rat | Mouse/Human |
| CD45 | 30-F11 | 103132 | Biolegend | PerCP/Cy5.5 | Rat | Mouse |
| CD45.1 | A20 | 110716 | Biolegend | APC-Cy7 | Mouse | Mouse |
| CD45.2 | 104 | 109828 | Biolegend | PerCP/Cy5.5 | Mouse | Mouse |
| CD62L | MEL-14 | 564108 | BD Bioscience | BV650 | Rat | Mouse |
| CD64 | X54-5/7.1 | 139323 | Biolegend | BV605 | Mouse | Mouse |
| CD90.2 (Thy1.2) | 53-2.1 | 140318 | Biolegend | BV605 | Rat | Mouse |
| CD103 | 2-E7 | 121432 | Biolegend | APC-Cy7 | Hamster | Mouse |
| CD127 | A7R34 | 135023 | Biolegend | BV421 | Rat | Mouse |
| Siglec-F (CD170) | S17007L | 155505 | Biolegend | PE | Rat | Mouse |
| IFN gamma | XMG1.2 | 25-7311-82 | Invitrogen | PECy7 | Rat | Mouse |
| IFN gamma | XMG1.2 | 505850 | Biolegend | APC-Cy7 | Rat | Mouse |
| IL2 | JES6-5H4 | 503814 | Biolegend | AF647 | Rat | Mouse |
| Live/Dead Fixable Stain |  | L34957 | ThermoFisher | Aqua | Rat | Mouse |
| Ly6C | HK1.4 | 128014 | Biolegend | PacBlue | Rat | Mouse |
| Ly6G | 1A8 | 127643 | Biolegend | BV711 | Rat | Mouse |
| MHCII (I-A/I-E) | M5/114.15.2 | 107622 | Biolegend | AF700 | Rat | Mouse |
| NK 1.1 | PK136 | 108718 | Biolegend | AF488 | Mouse | Mouse |
| Nur77 | 12.14 | 53-5965-82 | Invitrogen | AF488 | Mouse | Mouse |
| PD1 (CD279) | 29F.1A12 | 135216 | Biolegend | PE-Cy7 | Rat | Mouse |
| TNF alpha | MP6-XT22 | 506318 | Biolegend | PacBlue | Rat | Mouse |

**Supplemental table 2.** Lung Injury Scoring System from Aeffner et al. *Toxicol Pathol.* 2015.

| Parameter | Score Per High-Powered Field |  |  |
| --- | --- | --- | --- |
|  | 0 | 1 | 2 |
| A) Neutrophils in the alveolar space | None | 1-5 | > 5 |
| B) Neutrophils in the interstitial space | None | 1-5 | > 5 |
| C) Proteinaceous debris in the alveolar space | None | 1 | > 1 |
| D) Alveolar septal thickening | < 2 x normal | 2-4 x normal | > 4 x normal |

**Supplemental Figure 1.**
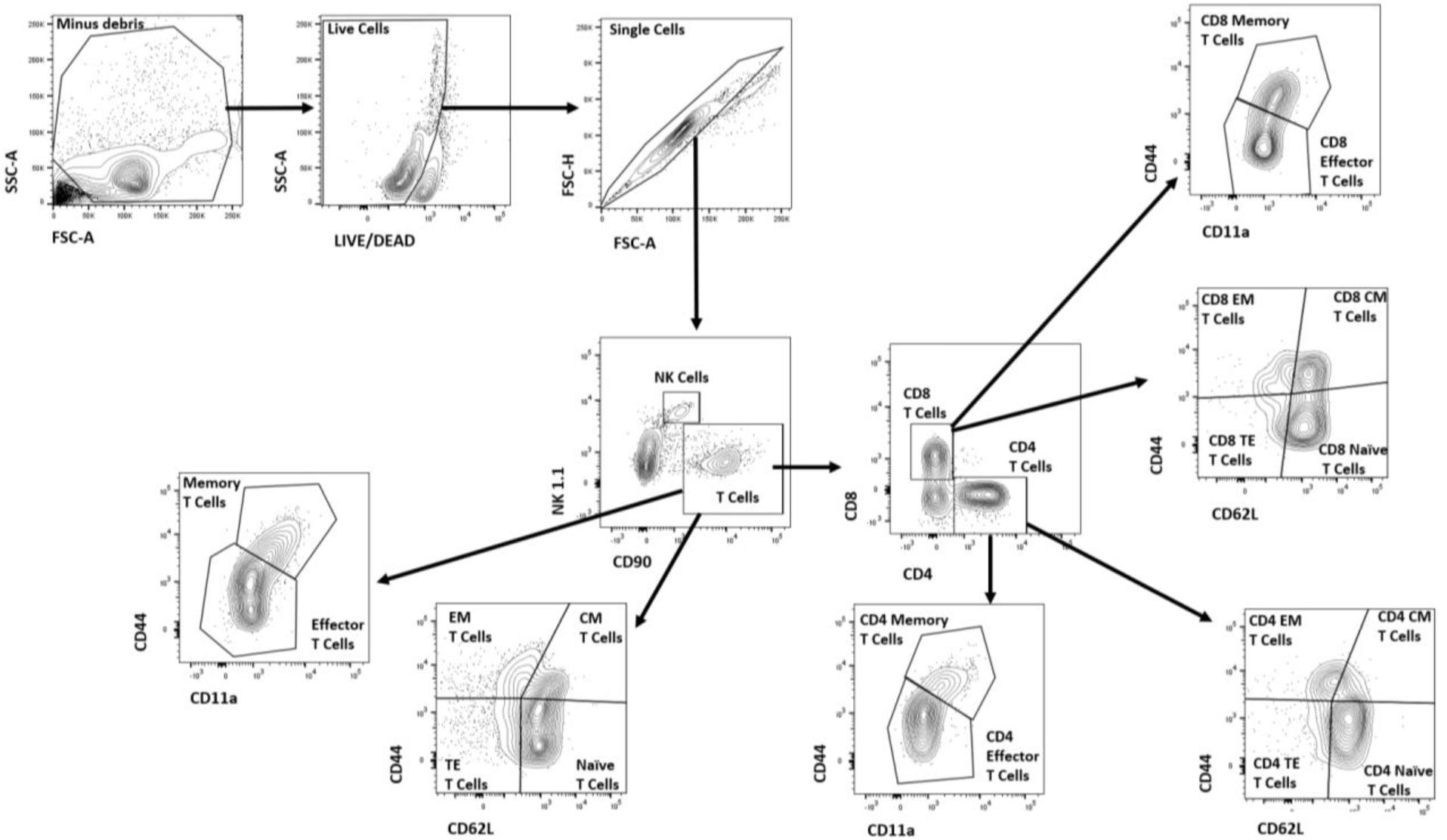
Gating strategy used to identify T cell subsets demonstrated in an Immune-Educated mouse at baseline. After exclusion of debris, dead cells, and doublets, T cells and NK cells were identified using CD90 and NK 1.1. CD90+ T cells were classified as Memory T cells (CD44+/CD11a+), Effector T cells (CD44-/CD11a-), Effector Memory (EM) T cells (CD44+/CD62L-), Central Memory (CM) T cells (CD44+/CD62L+), Naïve T cells (CD44-/CD62L+), Terminal Effector (TE) T cells (CD44-/CD62L-), CD8 T cells (CD8+), and CD4 T cells (CD4+). CD8 and CD4 T cells were further categorized as CD8+ or CD4+ Memory T cells, Effector T cells, EM T cells, CM T cells, Naïve T cells, and TE T cells.

